# p53 Controls Murine Gammaherpesvirus Latency and Prevents Infection-Associated *IgH/c-Myc* Translocations

**DOI:** 10.1101/2020.08.02.233148

**Authors:** Shana M. Owens, Jeffrey M. Sifford, Gang Li, Eduardo Salinas, Debopam Ghosh, Andrew D. Miller, Jason Stumhofer, J. Craig Forrest

## Abstract

Gammaherpesviruses (GHVs) establish life-long infections and cause cancer in humans and other animals. To facilitate chronic infection, GHV oncoproteins promote cellular proliferation and differentiation. Aberrant cell-cycle progression driven by viral oncogenes should trigger activation of tumor suppressor p53, unless p53 is functionally deactivated during GHV latency establishment. However, interactions of GHVs with the p53 pathway during the establishment and maintenance of latent infection are poorly defined. Here we demonstrate *in vivo* that p53 is induced specifically in infected cells during latency establishment by murine gammaherpesvirus 68 (MHV68). In the absence of p53, MHV68 latency establishment was significantly increased, especially in germinal center B cells, and correlated with enhanced cellular proliferation. However, enhanced latency was not sustainable, and MHV68 exhibited a defect in long-term latency maintenance in p53-deficient mice. Moreover, *IgH/c-Myc* translocations were readily detected in B cells from infected p53-null mice indicating virus-driven genomic instability. These data demonstrate that p53 intrinsically restricts MHV68 latency establishment and reveal a paradigm in which a host restriction factor provides a long-term benefit to a chronic pathogen by limiting infection-associated damage.

## INTRODUCTION

Gammaherpesviruses (GHVs) are DNA tumor viruses that include the human pathogens Epstein-Barr virus (EBV) and Kaposi sarcoma-associated herpesvirus (KSHV) and the rodent pathogen murine gammaherpesvirus 68 (MHV68), among others. Following a period of productive viral replication in various host tissues, GHVs establish lifelong chronic infections. This phase of the infectious cycle, referred to as latency, is preferentially maintained in B cells. The lifelong infections caused by GHVs place the host at risk for numerous cancers, including Burkitt lymphoma, Hodgkin lymphoma, primary effusion lymphoma, multicentric Castleman disease, and many others. Moreover, GHV-related malignancies represent serious complications for AIDS patients and transplant recipients. However, the precise mechanisms by which GHVs promote oncogenesis are incompletely defined.

In order to establish and maintain latency, GHVs stimulate and usurp normal B cell differentiation processes. This is accomplished through the actions of viral noncoding RNAs and oncoproteins, which, upon infection of naïve B cells, promote cellular activation reminiscent of a germinal center (GC) reaction. This process is thought to facilitate long-term latency in isotype-switched memory B cells. This model of GHV latency establishment is largely based on EBV infection and immortalization of primary B cells in culture and is supported by kinetic evaluations of cell-types that harbor MHV68 *in vivo* following experimental infection of mice [reviewed in (1, 2)]. However, normal somatic cells are remarkably sensitive to perturbations in cell cycle and differentiation status and respond to expression of viral and cellular oncogenes that drive aberrant cellular proliferation with potent tumor suppressor responses that arrest cellular growth and differentiation (3-5). To this point, EBV, KSHV, and MHV68 each encode viral proteins capable of inducing cell-cycle entry and transformation (6-9), but none of these viruses efficiently transforms infected cells.

For example, although most if not all cells are efficiently infected and begin to proliferate during the generation of lymphoblastoid cell lines (LCLs) following EBV infection of primary B cells in culture, only a small percentage of infected cells become immortalized and grow out (10, 11). EBV induces a period of cellular hyper-proliferation during LCL formation that triggers the activation of ataxia telangiectasia mutated (ATM) and checkpoint kinase 2 (Chk2), enzymes that function to inhibit further proliferation and subsequent immortalization (10). Similarly, KSHV triggers ATM activation and proliferative arrest upon infection of primary endothelial cells (12), and, although KSHV is capable of infecting and inducing proliferation of primary tonsillar B cells in culture, immortalization and long-term latent infection do not follow (13). Like EBV, MHV68 can immortalize primary cells in culture, but again the process is inefficient, requiring weeks of culture prior to the outgrowth of transformed cells (14).

As an intriguing *in vivo* correlate to these tissue culture studies, MHV68 induces potent lymphocyte proliferation during an infectious-mononucleosis (IM)-like syndrome following experimental infection of mice (1, 2). During this time, MHV68 is readily found in proliferating cells, and latently-infected cells reach a high point of ca. 0.5-1% of the total B cell population in the spleen (2). This is remarkably similar to EBV burdens in peripheral blood mononuclear cells during IM (15). However, for both MHV68 and EBV the percentage of latently infected cells rapidly declines to the steady-state levels present in long-term latent infection (1, 16). The contraction in latently infected cells correlates with maturation of the adaptive immune response (1, 2), but it also agrees with the notion that an intrinsic cellular defense program becomes activated to evoke apoptosis or cell-cycle arrest in a percentage of infected cells. This idea also is consistent with ATM and/or Chk2 enforcing barriers to LCL formation (10).

Whether innate restrictions present during GHV infection of primary B cells in culture accurately represent events that occur during natural infection of a host is not known. A role for ATM in restricting MHV68 infection *in vivo* recently was tested; however, ATM deficiency did not promote enhanced latent infection by MHV68 (17). Rather, ATM facilitates B cell latency (18, 19). This finding indicates that ATM does not impose a critical barrier to latent MHV68 infection *in vivo*.

Another candidate host protein for intrinsic latency restriction is tumor suppressor p53. p53 is considered a master regulator of genetic integrity, a postulate emphasized by the presence of inactivating p53 mutations in approximately half of all human cancers (20). Though constitutively expressed, p53 under normal physiologic cellular conditions is constantly targeted for proteolytic degradation (21). In response to potentially genotoxic cellular stresses, including DNA damage, oncogene expression, and viral infection, p53 becomes stabilized and activated through a variety of post-translational modifications (21). Active p53 regulates transcription of a number of cellular genes involved in cell-cycle arrest, DNA repair, and apoptosis (22). Interestingly, p53 is stabilized and p53-responsive transcripts are induced during the immortalization of primary B cells by EBV (23), and the KSHV-encoded cyclin D ortholog is sufficient in-and-of-itself to induce p53 (24). Exogenous induction of p53 reduces LCL formation by EBV, and conversely p53 inhibition facilitates endothelial cell proliferation following KSHV infection (12, 25). Although both EBV and KSHV encode latency proteins demonstrated to inhibit p53 in various biochemical evaluations (26, 27), established cell lines in which these viral inhibitors of p53 are highly expressed remain responsive to p53 agonists (23, 28), suggesting that these proteins inefficiently suppress p53 functions in latently infected cells. Moreover, it seems counterintuitive that a virus that establishes a life-long chronic infection would universally inhibit a tumor suppressor as critically important as p53. However, p53 mutations do occur frequently in endemic Burkitt lymphoma, a cancer characterized by stable EBV infection and the presence of *IgH/c-Myc* chromosomal translocations, which suggests that p53 functions limit GHV-related cancers *in vivo* (16). Whether p53 limits GHV latency and genomic instability has not been directly tested using *in vivo* models of pathogenesis.

Here we describe experiments that test the hypothesis that p53 is activated during GHV infection to restrict latency establishment and virus-driven cellular proliferation. Using MHV68 infection of WT and p53-deficient mice, we evaluate the functions of p53 in controlling latency establishment and maintenance by a GHV *in vivo*. Through this work we determine the impact of p53 on the infectious cycle of MHV68 and define the consequences of p53 deficiency for a GHV- infected host.

## RESULTS

### MHV68 induces p53 during latency establishment

p53 is activated in response to aberrant cellular proliferation and genotoxic stress, especially when driven by viral oncogene expression (29). To efficiently establish latency, GHVs encode proteins that promote cellular proliferation and differentiation (30), and primary cells infected with EBV or KSHV exhibit p53 induction (23). Whether p53 becomes stabilized and active during GHV latency establishment in a natural host is not known. To address this question, we infected mice intranasally with YFP-encoding MHV68 to mark infected cells (31) and evaluated p53 induction by flow cytometry during the establishment of latency in the spleen. On day 16 post-infection, a time point at which acute viral replication has waned, latent viral burdens in the spleen are at their peak, and the majority of infected B cells are proliferating and exhibit a germinal center phenotype (32, 33), levels of p53 in YFP^+^ cells were significantly higher than levels detected in mock-infected animals or YFP^-^ cells in spleens of infected mice (**Fig. 1A** and **1B**). Quantitative RT-PCR performed on B cells isolated from mock-infected or infected animals demonstrated increased transcription during infection of p53-responsive genes CDKN1A and MDM2 (**Fig. 1C**).

**Figure 1.**
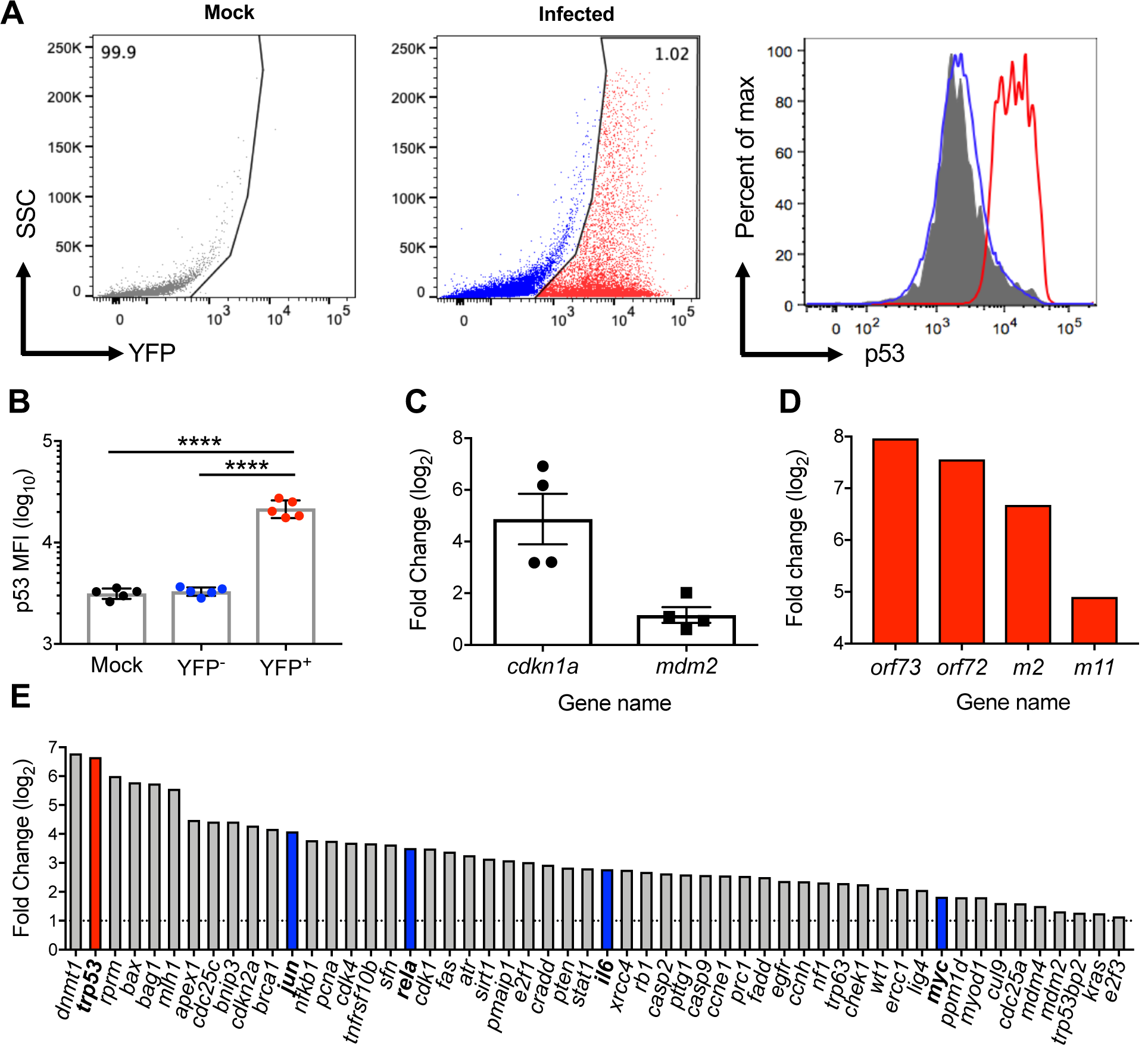
p53 is induced and active in latently infected splenocytes. (A and B) p53 is stabilized in MHV68-infected splenocytes. Flow cytometry was performed to evaluate p53 induction in spleen cells on day 16 after mock infection or infection of C57BL/6 mice with 10^4^ PFU of H2B-YFP MHV68. The mean fluorescence intensity of p53 staining in cells from mock-infected mice (gray), or YFP- (blue) and YFP+ cells (red) from infected animals are enumerated in B. Results are expressed as means +/- SEM. Mann-Whitney unpaired t test, **** p<0.0001. (C) Induction of p53 regulated transcription in MHV68-infected spleens. Quantitative RT-PCR analysis was performed on unsorted splenocyte RNA to detect *cdkn1a, mdm2*, and *β-actin* on day 16 post-infection. Data are relative transcript levels normalized to *β-actin* transcript abundance and compared to uninfected splenocytes by the Δ*C*_*T*_ method. (D-E) p53-related transcripts are increased specifically in infected cells. ZsGreen Ai6 reporter mice were infected intranasally with 10^4^ PFU of MHV68-Cre. ZsGreen^+^ or ZsGreen^-^ cells were sorted on day 16 post-infection. Relative viral (D) or cellular (E) transcript levels in ZsGreen^+^ cells normalized to *β-actin* were compared to ZsGreen^-^ splenocytes by the Δ*C*_*T*_ method. The *trp53* transcript is highlighted in red. Transcripts previously evaluated in MHV68 infection are denote with blue bars. The sorting strategy and enrichment of MHV68-infected cells is shown in Supplemental Figure 1.

To determine if p53 was transcriptionally active in infected cells, we infected Ai6 ZsGreen reporter mice (34) with a recombinant MHV68 virus that expresses Cre recombinase to induce ZsGreen expression (32). ZsGreen^+^ cells were sorted, and infection was confirmed by LD-PCR (**Fig. S1**). Viral latency transcripts were significantly enriched in ZsGreen^+^ cells relative to ZsGreen^-^ cells, and numerous p53 pathway-associated transcripts also were upregulated in comparative quantitative RT-PCR analyses (**Fig. 1D** and **1E**). Of note, TRP53 expression was ca. 100-fold higher in infected cells compared to ZsGreen^-^ cells. This is consistent with detection of a p53-related gene expression profile in RNA-seq analyses of splenocytes during MHV68 latency (36). Together these data indicate that p53 is activated *in vivo* during MHV68 latency establishment. The observation that p53 stabilization occurred almost exclusively in YFP^+^ cells demonstrates that p53 activation is not a general consequence of viral infection, but represents a cell-intrinsic response to MHV68 infection.

### MHV68 latency establishment is enhanced in p53-deficient mice

Given that p53 induction was predominant in infected cells, we hypothesized that p53 is activated as an inherent host-cell response to limit MHV68 latency establishment. To test this hypothesis, we evaluated MHV68 infection in p53^+/+^ and p53^-/-^ mice. Although p53 is reportedly an interferon (IFN) stimulated gene that functions in limiting acute viral replication and MHV68 replication is controlled by type I IFNs (37, 38), viral loads were equivalent in lungs of p53^-/-^ and WT animals after intranasal inoculation and mortality was not observed during the acute phase of infection (**Fig. S2**). This indicates that p53 is not critical to control MHV68 during the acute phase of infection and is consistent with viral shut-down of p53 function during lytic replication (39, 40). In contrast, the frequency of cells harboring MHV68 genomes during latent colonization of the spleen on day 16 post-infection was 14-fold higher in p53^-/-^ animals than WT littermates, with approximately 1 in 20 p53^-/-^ cells infected vs. 1 in ca. 280 WT cells (**Fig. 2A** and **Table 1**). Similar enhancement of infection was observed in p53^-/-^ mice when cells were analyzed by flow cytometry to enumerate YFP^+^ cells (**Fig. 2B**). Moreover, immunohistochemical analyses demonstrated an increase relative to WT animals in the number of infected cells present within B cell follicles of p53-null mice (**Fig. 2D**).

**Table 1.**
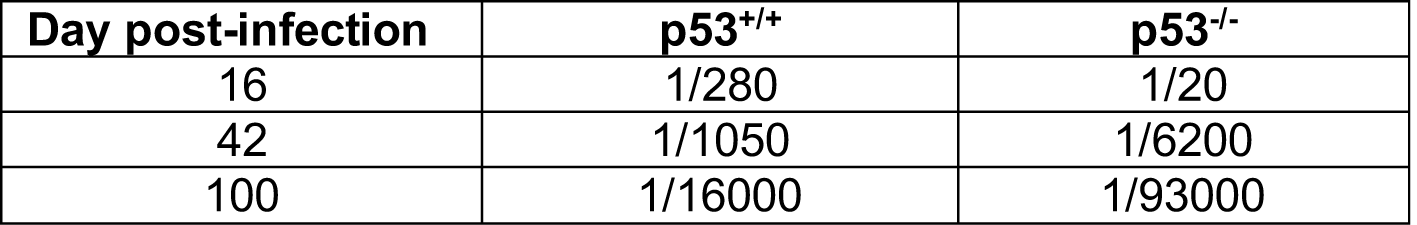
MHV68 genome loads in p53^+/+^ and p53^-/-^ mice.

**Figure 2.**
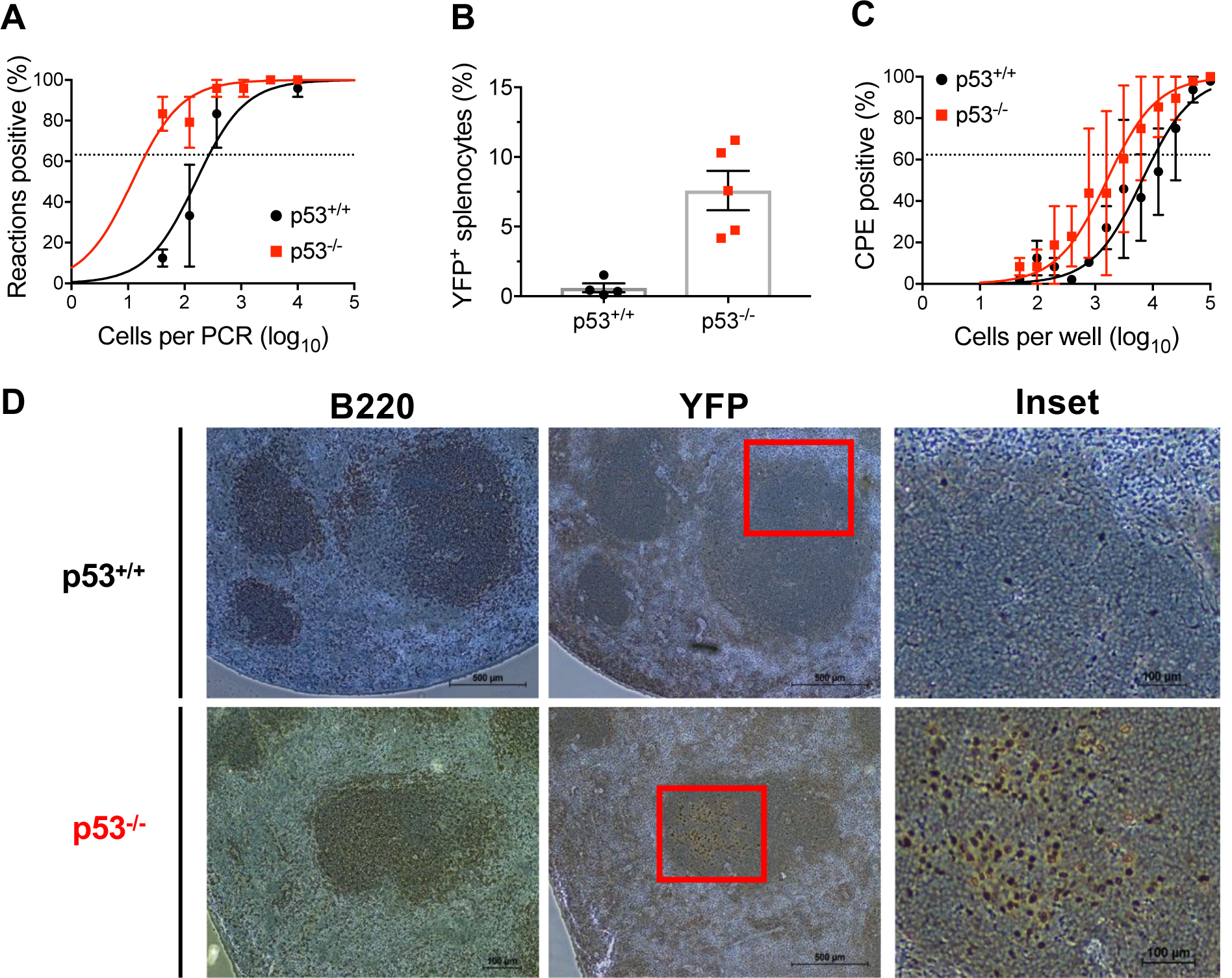
MHV68 latency establishment is enhanced in p53-deficient mice. (A and B) Increased detection of MHV68 infected cells in p53 knockout mice. p53^-/-^ or p53^+/+^ were infected with 10^4^ PFU of H2B-YFP MHV68. Limiting-dilution PCR to detect viral genomes (A) or flow cytometry to detect YFP (B) was performed on splenocytes on day 16 post-infection. Results in B are means +/- SEM. Mann- Whitney unpaired t test, ** p<0.005. (C) Reactivation from latency is normal in the absence of p53. Splenocytes were cultured ex vivo on an indicator MEF monolayer to evaluate MHV68 reactivation capacity from p53^-/-^ or p53^+/+^ mice. Acute replication and adaptive immune responses were normal in p53^-/-^ mice (Supplemental Figures 2 and 3). (D) MHV68 infection of splenic follicles is enhanced in p53 knockout mice. Spleens were harvest on day 16 after infection, and immunohistochemical analyses were performed to detect splenic follicle B cells (B220) and MHV68-infected cells (YFP). Representative images are shown. Scale bars indicate 500 μm or 100 μm for the inset.

A percentage of cells latently infected with MHV68 undergo spontaneous reactivation when cultured ex vivo, and defects in innate immune control can promote enhanced reactivation (41-43). While p53^-/-^ animals contained more reactivation-competent splenocytes than WT mice (**Fig. 2C**), the increase directly correlated with the increase in the number of cells harboring MHV68. The reactivation efficiency therefore was not directly influenced by p53 status (**Table 1**). We also did not detect in these assays an increase in preformed infectious virus, which is indicative of ongoing persistent viral replication (44, 45), in p53^-/-^ mice (**Fig. 2C**). These findings demonstrate that the absence of p53 does not result in persistent viral replication or hyper- reactivation. Since there is not a reactivation defect, it also is unlikely that increased latency occurs as a consequence of failed reactivation.

Finally, MHV68-specific adaptive immune responses occurred normally in p53^-/-^ mice, as virus-specific IgG in serum, T cell activation, and induction of MHV68 antigen-specific CD8^+^ T cells were similar in both WT and p53-null animals infected with MHV68 (**Fig. S3** and **S7**). Together, these data demonstrate that the absence of p53 correlates with enhanced MHV68 latency establishment and suggest that p53 promotes intrinsic cellular resistance to chronic MHV68 infection.

### p53 controls cellular proliferation and death during MHV68 latency establishment

To gain insight into roles for p53 in controlling MHV68 latency establishment, we evaluated how the presence or absence of p53 influenced the infection-associated expansion of specific cell types and the impact of p53 on MHV68 targeting of distinct cell populations. During latency establishment, cellularity in spleens increases as a correlate of an IM-like syndrome induced by MHV68 (46). Although spleens from mock-infected p53^-/-^ and WT mice contained equivalent numbers of cells, following infection p53^-/-^ spleens exhibited a significantly greater increase in cellularity compared to WT mice (**Fig. 3A** and **S7**). Quantification of specific lymphocyte populations by flow cytometry revealed enhanced expansion CD19^+^B220^+^ B cells in p53^-/-^ mice (**Fig. 3B**), while T and NK cell numbers were similar between WT and knockout animals (**Fig. S4**). For particular B cell subsets, the number of GC B cells was significantly increased (**Fig. 3C**). Plasmablasts also were present in greater numbers, but did not achieve significance (**Fig. 3D**). Moreover, total B cells, GC B cells, and plasmablasts that harbored MHV68 were increased in the absence of p53 (**Fig. 3E-H**).

**Figure 3.**
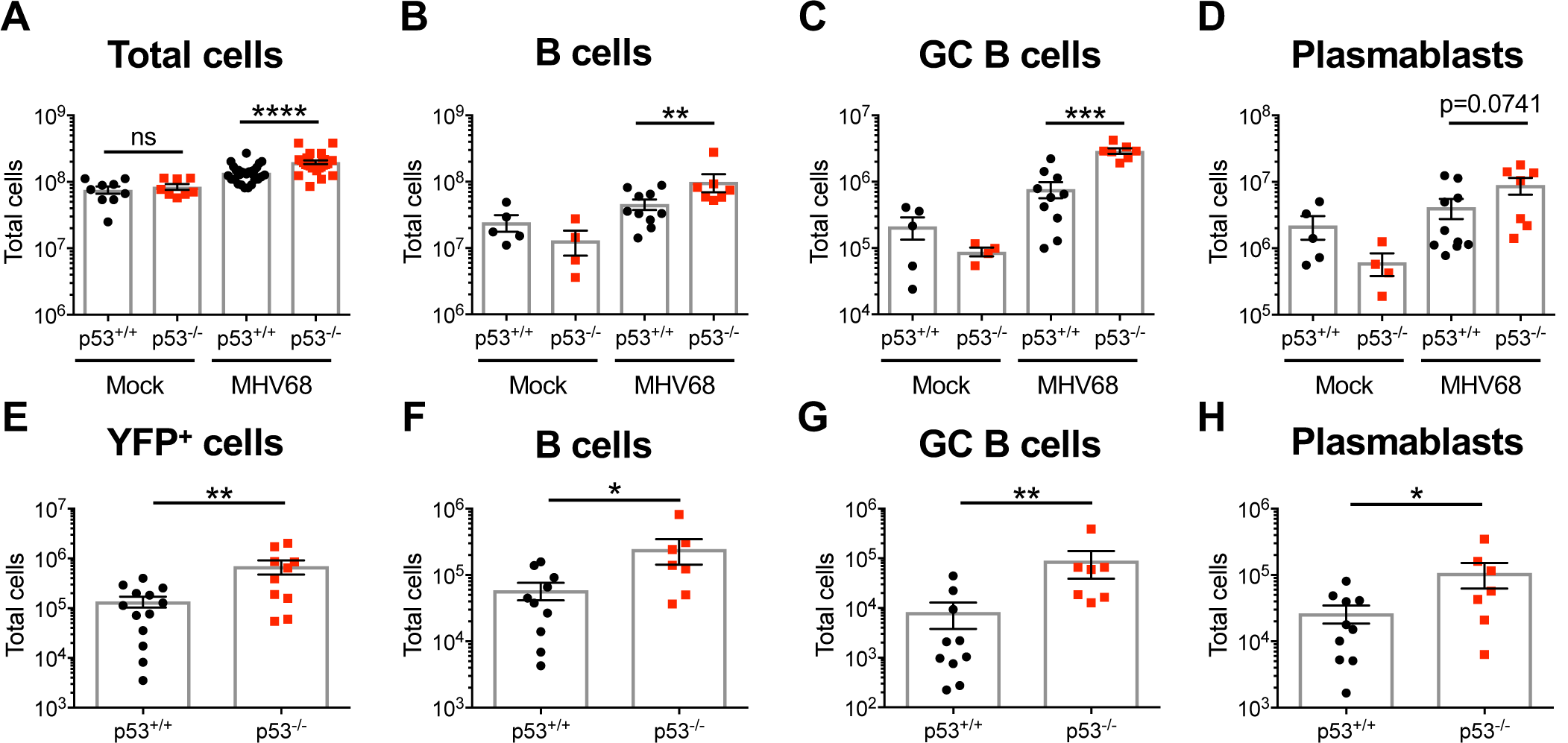
B cell expansion and infection is increased in p53-deficient mice. (A-D) The infection-associated increase in spleen cell numbers is enhanced in the absence of p53. On day 16 after mock infection or infection with H2B-YFP MHV68, total cells in spleens from p53^+/+^ or p53^-/-^ were quantified and analyzed by flow cytometry. B cells were gated as CD19^+^/B220^+^, GC B cells as GL7^+^/CD38^lo^, and plasmablasts as CD138^+^/B220^lo^. Numbers of other lymphocyte populations were equivalent in p53^-/-^ and p53^+/+^ spleens (Supplemental Figure 4). Gating strategies are shown in Supplemental Figure 7. (E-H) Total numbers of specific B cell subsets infected by MHV68 is increased in the absence of p53. Data represent 2-3 independent infections with a minimum of 3 mice per group. Results are expressed as means +/- SEM. Mann-Whitney unpaired t test * p<0.05, ** p<0.005, ***p<0.0005, **** p<0.0001.

As a control to determine whether the increase in B cell numbers in p53^-/-^ mice was a general response to infection, we infected knockout mice with mLANA-null MHV68, a mutant virus that undergoes acute replication and triggers adaptive immunity, but fails to disseminate to the spleen following intranasal inoculation (47-49). In contrast to WT MHV68, mLANA-null virus did not promote a similar increase in splenocyte numbers after infection of p53^-/-^ mice (**Fig. S5**). Together, these findings demonstrate that p53 controls both the expansion of B lineage cells and their infection by MHV68. These data also suggest that the expansion of spleen cells in p53^-/-^ mice is driven by the presence of MHV68 and is not simply a consequence of the general immune response to viral infection.

Given that GHVs drive B cell proliferation in order to establish and maintain latency (30, 50) and p53 restricts cellular proliferation or promotes apoptosis downstream of viral oncogene expression (51-53), we hypothesized that p53 functions to limit MHV68-driven cellular proliferation and/or induce death in infected cells. Using *in vivo* EdU-incorporation assays to mark cells actively replicating their DNA, we detected similar numbers of EdU^+^ cells in both p53^-/-^ and WT animals that were mock infected, indicating equivalent basal cellular proliferation rates. Infection with MHV68 led to an increase in the number of EdU^+^ GC B cells irrespective of genotype; however, spleens of p53-deficient mice contained more EdU^+^ GC B cells than WT mice (**Fig. 4A** and **S8A**). Cell death was only modestly reduced in the absence of p53 in evaluations using annexin V and propidium iodide staining of infected splenocytes (**Fig. 4B** and **S8B**). These data suggest that a mechanism by which p53 limits MHV68 latency establishment is through restriction of cellular proliferation. The enhanced expansion and infection of GC B cells in p53^-/-^ mice is consistent with this interpretation and suggests that the highly proliferative GC B cells are the main cell type in which p53-mediated restriction occurs.

**Figure 4.**
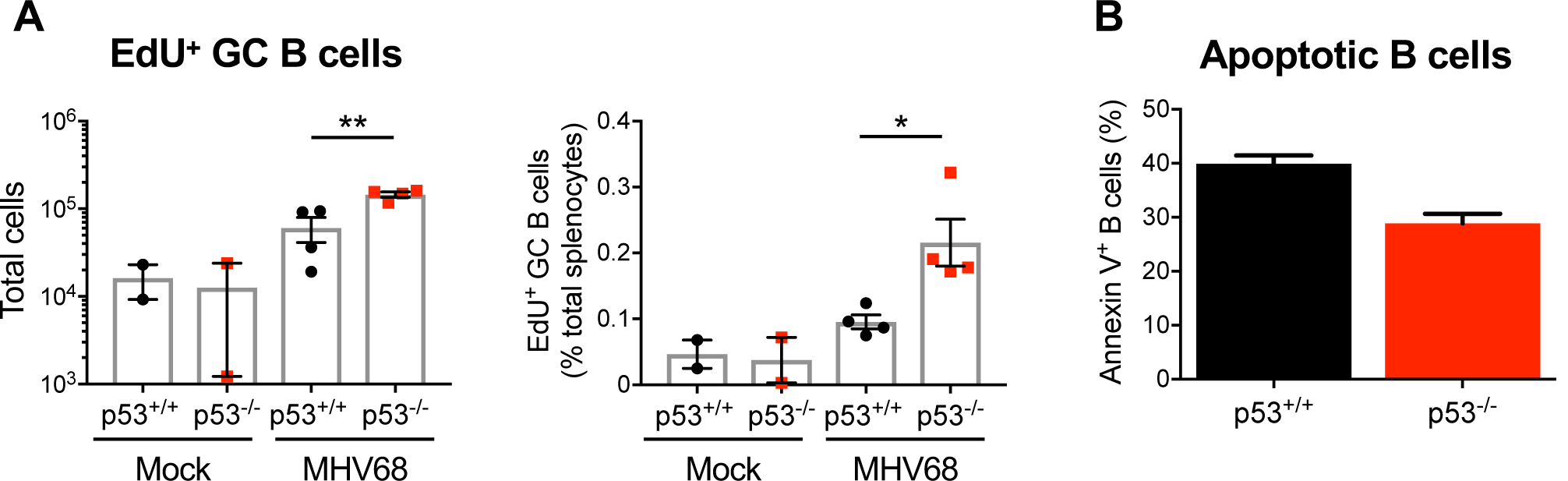
p53 controls cellular proliferation during MHV68 latency establishment. (A) More cells are undergoing DNA replication in infected p53 knockout mice. Mock-infected and MHV68- infected mice were injected intraperitoneally with EdU on day 16 post-infection. Two hours after injection, spleen cells with EdU incorporation in newly synthesized DNA were labeled by Click chemistry. EdU+ germinal center B cells were detected by flow cytometry (B cells were gated as CD19^+^/B220^+^, GC B cells as GL7^+^/CD38^lo^). Results are means +/- SEM. Mann-Whitney unpaired t test * p<0.05, ** p<0.005 (B) B cell apoptosis is slightly reduced in the absence of p53. On day 16 post-infection, apoptotic B cells (CD19^+^, Annexin V^+^, and/or PI^+^) in infected p53^-/-^ or p53^+/+^ mice were evaluated by flow cytometry. splenocytes were harvested. Splenocytes were stained for CD19, Annexin V, and propidium iodide. Results are means +/- SEM. Gating strategies are shown in Supplemental Figure 8.

### Long-term MHV68 latency is reduced in p53-deficient mice

After the initial virus-driven B cell activation and expansion associated with MHV68 infection, both total cell numbers and the frequency of cells harboring viral genomes contract to reduced levels that are maintained during long-term infection. To determine the impact of p53 on long-term MHV68 latency, we evaluated latency at later times during infection. In contrast to the increased latency observed on day 16 post-infection, the frequency of cells harboring MHV68 in spleens of p53^-/-^ mice were reduced compared to WT mice on day 42 post-infection (**Fig. 5A**). Total cells, total B cells, GC B cells, and plasmablasts were present in spleens at equivalent numbers for knockout and WT animals at this time point (**Fig. 5C-F**). Thus, the increase in cell numbers and latency observed in the absence of p53 during colonization of the host was not sustained long-term. These findings demonstrate that p53 is not critical for limiting continued cellular and viral expansion after MHV68 latency is established. The reduction in latency observed, a phenotype that was maintained up to 100 days post-infection (**Fig. 5B**), suggests that, despite restricting initial latent colonization of the host, p53 ultimately plays a pro-viral role in long-term, chronic infection by MHV68.

**Figure 5.**
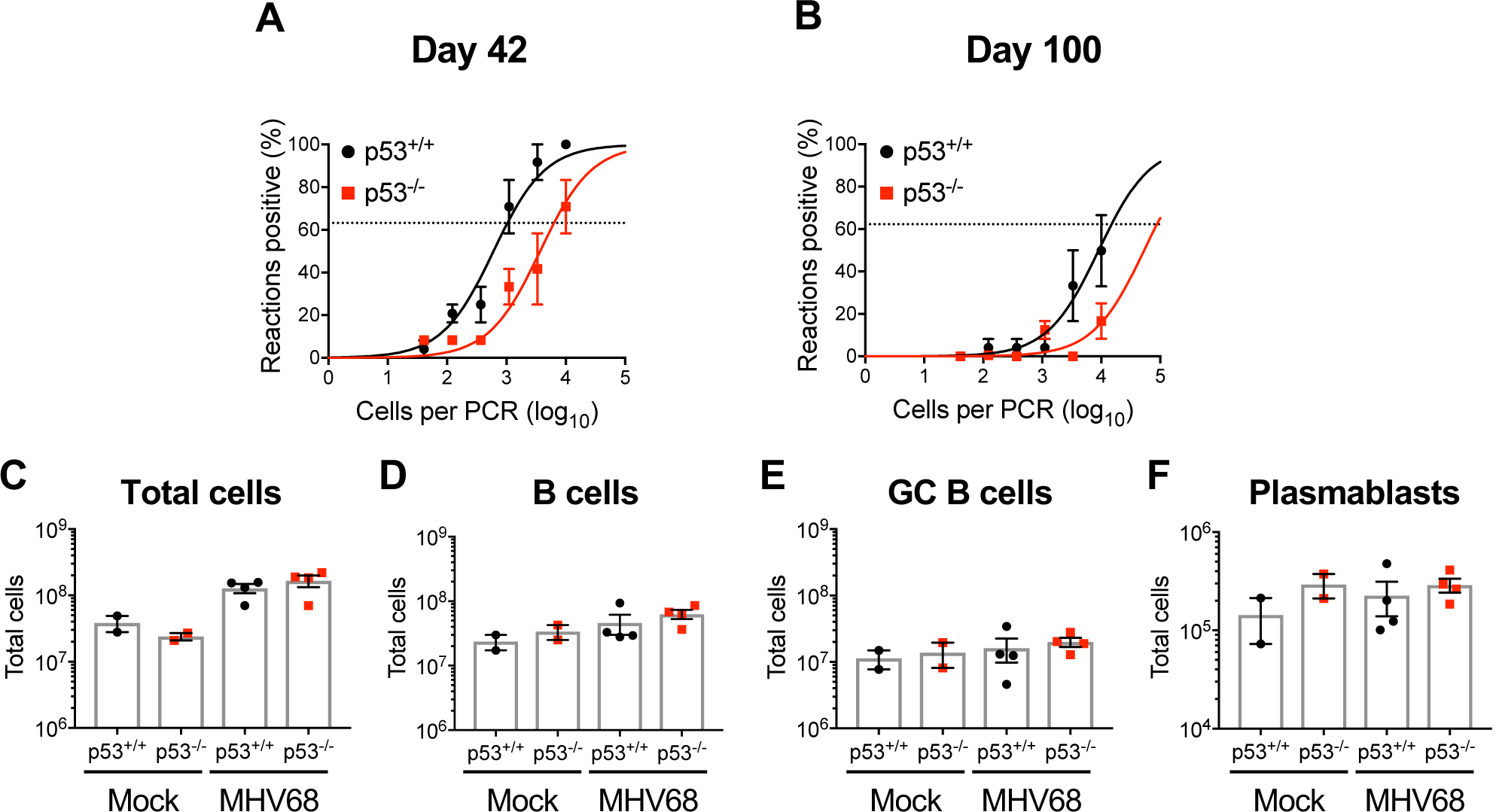
MHV68 latency is attenuated in long-term infected p53-deficient mice. (A-B) Reduction in MHV68-positive cells at late timepoints in the absence of p53. On day 42 (A) or 100 (B) after infection with MHV68, a limiting-dilution PCR analysis was performed to determine the number of cells maintaining MHV68 genomes. (C-F) Numbers of specific cell types in spleens are equivalent in WT and KO mice on day 42 post-infection. Total cells in spleens from mock-infected and MHV68-infected p53^+/+^ or p53^-/-^ were quantified and analyzed by flow cytometry. B cells were gated as CD19^+^/B220^+^, GC B cells as GL7^+^/CD38^lo^, and plasmablasts as CD138^+^/B220^lo^. Results are means +/- SEM. No significant differences were present by Mann-Whitney unpaired t test.

### *IgH/c-myc* translocations occur in p53-deficient mice infected with MHV68

Off-target DNA breaks caused by activation-induced cytidine deaminase (AID), the enzyme responsible for class-switch recombination and somatic hypermutation in activated B cells, can promote chromosomal translocations, especially in the absence of p53 (54, 55). Among the best characterized of these translocations is the *IgH/c-myc* translocation, which is also a hallmark of EBV-associated Burkitt lymphoma (56). Given that expansion and infection of GC B cells was increased in p53-deficient mice, we hypothesized that the absence of p53 correlates with accumulation of *IgH/c-myc* translocations during MHV68 infection.

To test this hypothesis, we performed an *IgH/c-myc* translocation-specific PCR with single-copy sensitivity (**Fig. S6**) on B cells isolated from spleens of mock-infected and infected mice. *IgH/c-myc* translocations were not detected in mock-infected animals or WT mice infected with MHV68. However, as early as 16 days post-infection *IgH/c-myc* translocations were detected in all MHV68-infected p53^-/-^ mice examined (**Fig. 6** and **S6**). Sequencing of the translocation- specific PCR products indicated that all rearrangements that amplified on day 16 post-infection represented unique translocation events (**Table 2**). For one infected animal evaluated 42 days post-infection, the same translocation junction was detected multiple times, which indicates clonal proliferation of a single translocation-positive cell. These data demonstrate that MHV68 infection promotes *IgH/c-myc* translocations in p53-deficient mice. These findings illustrate the importance of p53 in limiting mutations during infection by a GHV, especially chromosomal translocations such as those that occur in Burkitt lymphoma.

**Table 2.**
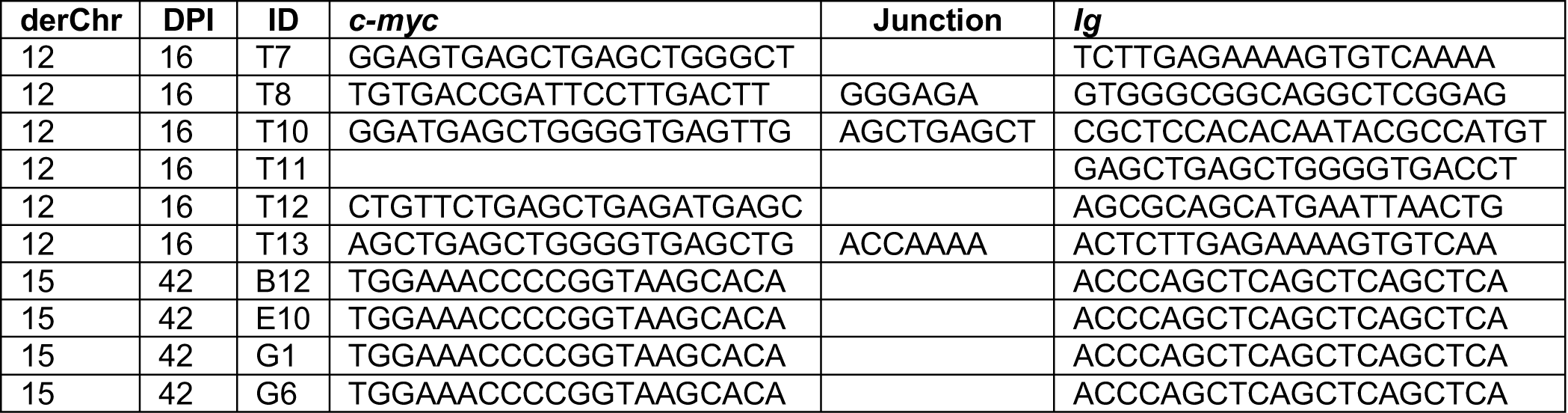
*IgH/c-myc* translocation sequences in p53^-/-^ mice.

**Figure 6.**
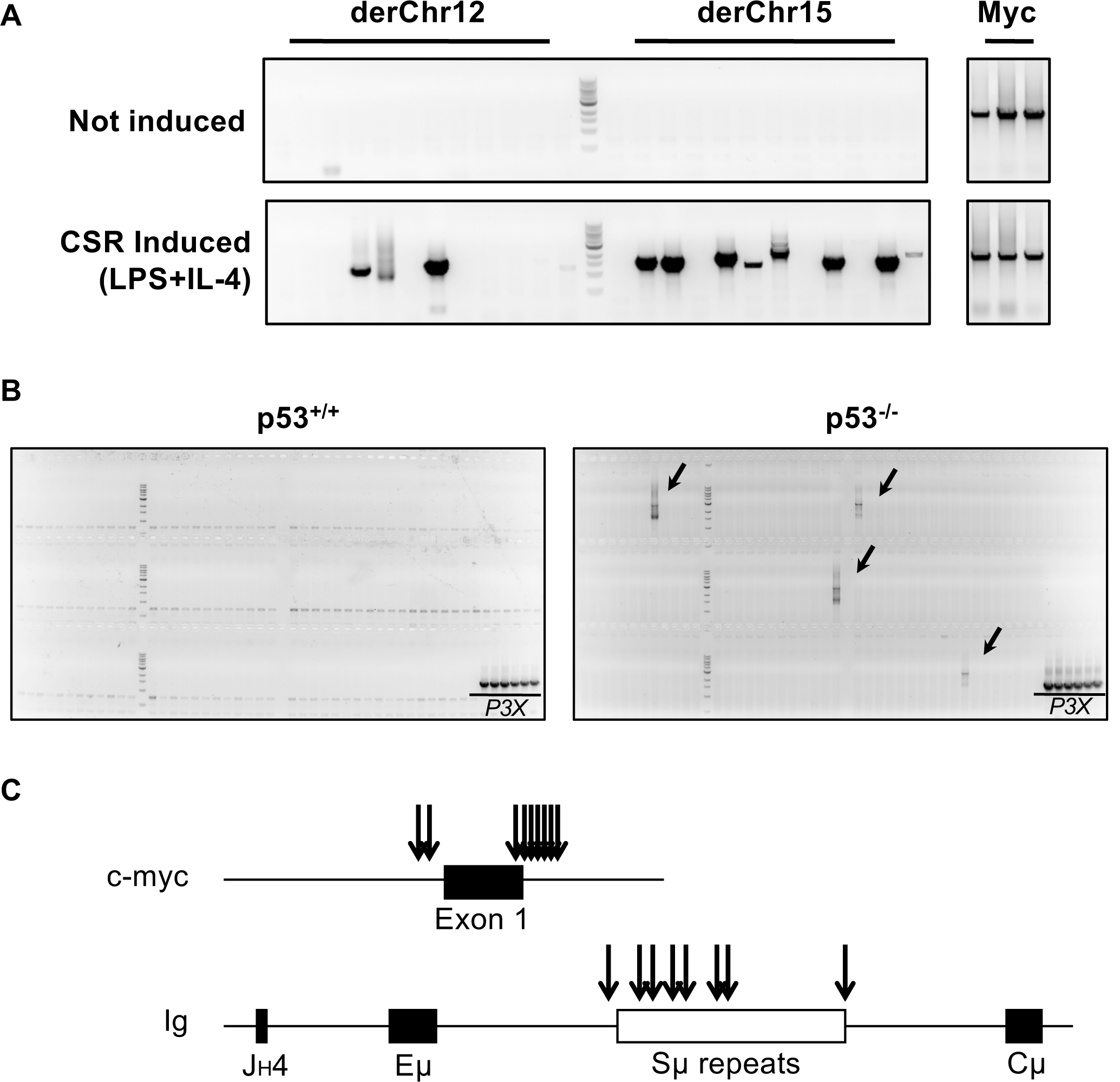
Infection of p53^-/-^ mice causes genomic instability. (A) Development of a high-sensitivity PCR approach to detect translocations. Primary p53^-/-^ B cells were isolated from naïve animals for *ex vivo* culture. B cells were treated with LPS and IL-4 to induce class-switch recombination (CSR) and chromosomal translocations. Each lane represents PCR using DNA from 10^5^ p53^-/-^ B cells. The apparent translocation frequency is 50 per 10^7^ cells, which is more sensitive than a previous PCR- based translocation analysis (frequency of 2 per 10^7^ cells, Ramiro et al, 2006). PCR sequencing confirmed that products were *IgH/c-myc* translocations. (B) Detection of *IgH/c-myc* tranlsocations after MHV68 infection of p53^-/-^ mice. Translocation PCR was performed on B cells isolated from either p53^+/+^ or p53^-/-^ mice 42 days post-infection. Positive bands (arrows) were excised and sequenced, confirming the amplification of *IgH/c-myc* translocations. Each lane represents PCR of DNA from 10^5^ B cells. DNA from P3X plasmacytoma cells served as a PCR positive control. No translocations have been detected in WT mice or mock-infected p53-/- mice. IgH/c-myc translocations also were detected on day 16 post-infection of p53-/- mice (Supplemental Figure 6). (C) Schematic representing breakpoint locations of sequenced fragments. Specific nucleotide sequences are available in Table 2.

**Figure 7.**
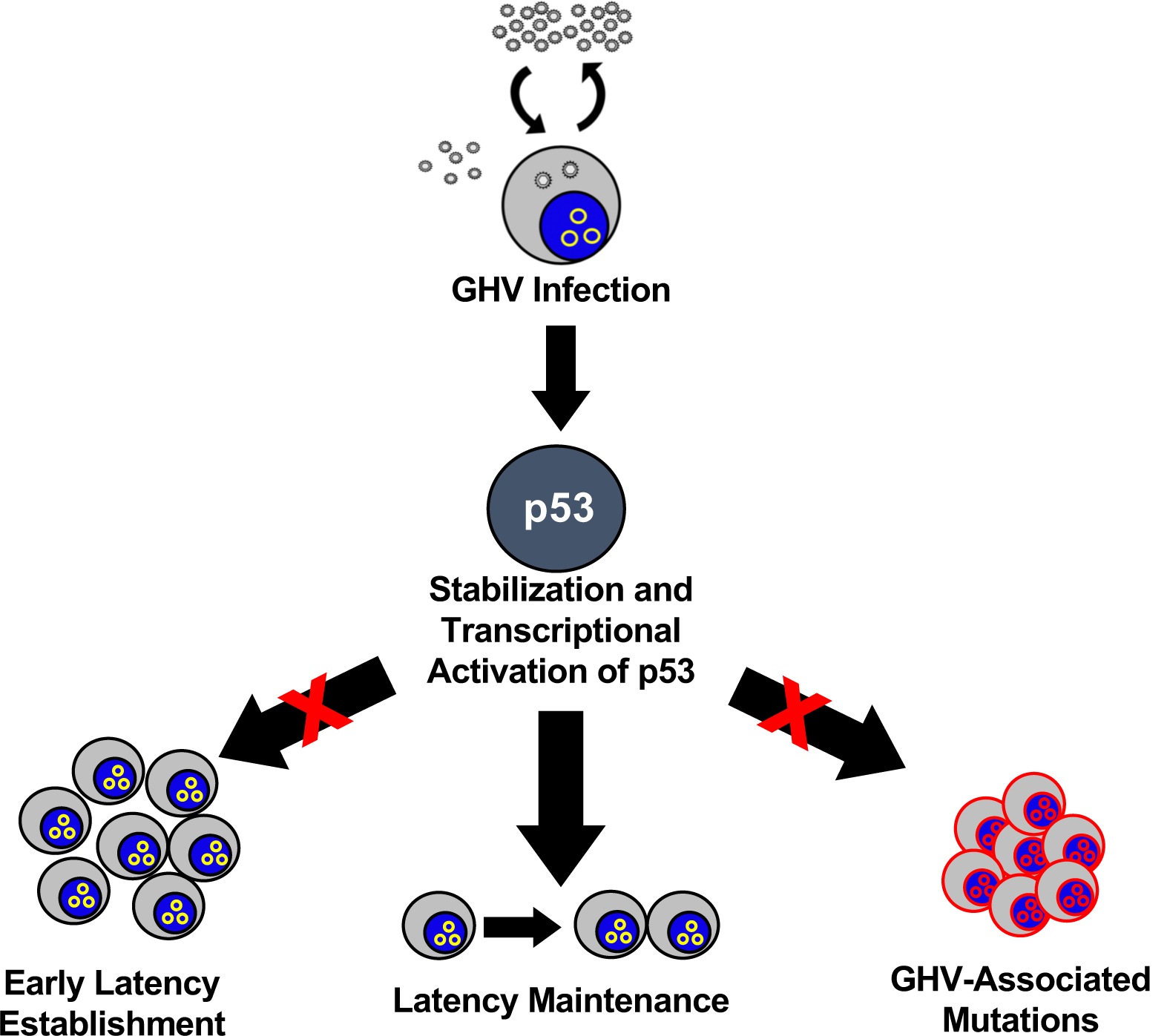
Model of p53 function during MHV68 latency establishment and maintenance. p53 is active during GHV infection and functions to restrict early latency establishment and prevent GHV-associated genomic instability, which promotes maintenance of latently infected cells.

## DISCUSSION

Data presented here demonstrate that p53 is induced specifically in infected cells during the establishment of latency by MHV68 in mice. Because early expansion of the latent virus pool was enhanced in the absence of p53 and correlated with increased GC B cell numbers and infection by MHV68, our data support a model in which virus-driven B cell proliferation leads to the activation of p53 (**Fig. 8**). In agreement with known responses to viral oncogene expression (51-53), p53 then serves to counteract infected cell expansion, while also limiting infection-related genomic instability. However, the absence of p53 was detrimental to long-term chronic infection, suggesting a paradigm in which a pathogen that has co-evolved with its host is fine-tuned to an intrinsic host response in order to limit pathology while maximizing latent viral persistence.

### p53 induction and impact on chronic infection

It is well established that GHVs induce and benefit from B cell proliferation and differentiation during colonization of the host (31, 50). This is achieved by the expression of the various latency genes that facilitate this process, and we suspect that augmentation of normal B cell differentiation pathways by GHV latency proteins is the primary trigger for p53 activation. EBV, which also causes p53 induction upon infection of primary human B cells (23), uses a coordinated latency gene expression program to similarly drive cellular proliferation and differentiation (8, 57). Both MHV68 and EBV induce and benefit from c-Myc during latency, and KSHV infected PEL cells are dependent on c-Myc activity for survival (58, 59). For MHV68, c-Myc is stabilized by mLANA in GC B cells (60). Since, aberrant c-Myc expression is a potent activator of p53 (61), the increased c-Myc activity driven by mLANA may promote p53. V-cyclin, a conserved viral ortholog of cellular cyclin D encoded by *ORF72* of MHV68 and KSHV, is another likely candidate to induce p53. When expressed as a transgene in mice, MHV68 v-cyclin promotes cell cycle progression and lymphomagenesis (9). Exogenous expression of KSHV v-cyclin in somatic cells promotes p53 induction with cell-cycle arrest and apoptosis, and v-cyclin drives tumorigenesis in p53^-/-^ mice (62). The MHV68 latency protein M2 is proposed to function in a manner analogous to EBV proteins LMP1 and LMP2a, potentiating MHV68 infection of GC cells, B cell survival, and plasma cell differentiation (63, 64). Although p53 normally is repressed by Bcl- 6 during GC reactions to enable rapid cellular proliferation and AID-directed mutagenesis of the Ig locus (65), we hypothesize that augmentation of the normal GC response by M2 might also engage p53. Since p53 restricts genomic instability caused by AID, it will be of interest to determine if MHV68-infected GC cells experience prolonged AID expression, increasing the likelihood of aberrant targeting and mutations. Consistent with this idea, EBV-infected tonsillar B cells express higher levels of AID than do non-infected cells (66, 67).

A common pathway for p53 activation during viral infection involves an ATM-mediated DNA damage response (DDR) (29). Infection of primary endothelial cells with KSHV leads to induction of an ATM-mediated DDR, a phenotype dependent on v-cyclin, that prevents the proliferation of infected cells (12). Infection of PBMCs with EBV likewise triggers an ATM-mediated DDR that must be overcome for LCL formation (10, 68). It is reasonable to suggest that MHV68 likewise triggers p53 through an ATM-mediated DDR, and we previously demonstrated that this is the case during lytic viral replication (39). Whether MHV68 causes ATM activation during latency establishment specifically in infected B cells has not been directly evaluated. However, the observation that B-cell-specific deletion of ATM correlates with a global defect in MHV68 latency (19), rather than the early enhancement in infection we observed in the absence of p53 during initial colonization of the host, suggests that ATM and p53 function independently in latently infected cells in vivo. Alternatively, it is possible that p53 induction occurs during MHV68 infection as a consequence of replication stress due to virus-driven cellular proliferation. Indeed, in addition to p53-related gene expression, RNAseq analyses of MHV68-infected splenocytes revealed enhanced transcription of genes involved in the ATM and Rad3-related (ATR) signaling pathway, a major responder to replication stress (36). Moreover, EBV also drives ATR induction during LCL formation (69, 70).

The finding that MHV68 latency establishment is enhanced in the absence of p53 in a manner that correlates with increased cellular proliferation and expansion of GC B cells is consistent with p53 functioning as an intrinsic restriction factor for MHV68 latency *in vivo*. These phenotypes also agree with MHV68 not blocking p53 function, or doing so only minimally, during latency. Whether GHVs inhibit p53 as a matter of course during latency has become controversial. Although KSHV LANA was demonstrated to potently inhibit p53 (26), several subsequent studies have challenged this original finding. PEL cells harboring latent KSHV and expressing high levels of LANA, as well as other latency-associated transcripts, remain sensitive to p53 agonists, responding to treatment with potent p53 stabilization, p53-related gene expression, and cell death (28, 71, 72). In contrast, KSHV-infected cells are resistant to p53-mediated death during lytic replication (73). This is also true for MHV68, in that LANA does not in-and-of-itself inhibit p53, but is required for p53 inhibition as the lytic cycle progresses (39, 40). In retrospect, since p53 inhibition is potently transforming, the assertion that p53 is not directly inhibited during long-term chronic infection agrees with the truth that GHV infection does not cause cancer in the majority of infected individuals.

GHVs co-evolved with their hosts, resulting in a virus-host relationship in which severe disease is not a common outcome. Considering that *IgH/c-myc* translocations were detected only in infected p53^-/-^ animals and not WT mice, our findings support a paradigm in which activation of p53 by MHV68 establishes a threshold in which virus-driven cellular proliferation is sufficient to permit colonization of the host, but insufficient to drive mutations and cancer. The deficit in long-term chronic infection indicates that the virus benefits from the restriction imposed by p53. As such, the long-term latency pool likely evolves from cells that do not accumulate DNA damage or are able to repair any potential genetic aberrancies. By extension, mutations in host cell genomes that normally would be limited by p53 function in WT mice ultimately lead to the loss of such cells from the long-term latent pool. The *IgH/c-myc* translocation PCR reveals a single chromosomal aberration, but the potential for numerous not-yet-tested genetic anomalies to be lurking within the genomes of infected p53-deficient cells is high. In essence, p53 is a cytoprotective molecule that functions to restrict cellular proliferation until various stresses are resolved or mutations are repaired, progressing to apoptosis as a last resort (3, 21). The fate of cells that have initiated p53 responses due to MHV68 infection is not clear from our work; it will be of interest to determine if such cells go on to become part of the latent pool during long-term chronic infection or contribute to latency by other means.

Based on the detection of *IgH/c-Myc* translocations in MHV68-infected p53-null mice, but not MHV68-infected WT mice or uninfected mice of either genotype, it is clear that latent infection has mutagenic potential. *IgH/c-myc* translocations are a hallmark of EBV-associated Burkitt lymphoma, and high EBV genome levels in individuals correlates with an elevated risk of Burkitt lymphoma development in endemic areas (56). We expect that increased viral loads are similarly a correlate of MHV68 disease in p53^-/-^ mice. Importantly, endemic Burkitt lymphoma also correlates with high incidence of co-infection by the malaria causing parasite, *Plasmodium falciparum*. It was recently demonstrated in mice that p53 prevents *IgH/c-Myc* translocations (and other genetic abnormalities) caused by prolonged GC responses during infection by *P. chabaudi*, a murine malaria parasite (74). Our work highlights the feasibility and potential benefit of combining these two models for co-infection studies aimed at elucidating the mechanisms by which GHV and *Plasmodium* infections synergize to promote the development of Burkitt lymphoma.

## MATERIALS AND METHODS

### Cell culture and viruses

Swiss albino 3T3 fibroblasts were purchased from ATCC. Murine embryonic fibroblasts (MEFs) were harvested from C57BL/6 mice embryos and immortalized as previously described (44). All cells were cultured in DMEM supplemented with 10% fetal bovine serum, 2 mM L-glutamine, and 100 U/ml penicillin/streptomycin. Cells were maintained at 37°C in 5% CO_2_. Viruses used in this study were previously described and include WT MHV68 (75), H2B-yellow fluorescent protein (YFP)-expressing MHV68 (76), and mLANA-null MHV68 (73.STOP; (48). Viral stocks were generated as previously described (40). Viral titers were determined by MHV68 plaque assay (77).

### Mice and infections

All mice were housed and cared for according to the guidelines of UAMS department of laboratory animal medicine and all state and federal requirements. p53-null mice on a C57BL/6 background (B6.129S2-*Trp53*^*tm1Tyj*^*/*J) were purchased from Jackson Laboratories and bred p53^+/-^ x p53^+/-^ to get a distribution of p53^+/+^, p53^+/-^, and p53^-/-^ progeny. 7-11 week-old mice were infected with 10^4^ PFU of WT MHV68, H2B-YFP MHV68, or mLANA-null MHV68 by intranasal inoculation or by intraperitoneal injection. Mice were sacrificed according to normal endpoint protocols at the first sign of illness or tumor development. Blood was collected on day 0, 16 and 42 post-infection via the submandibular vein.

### Splenocyte isolation and limiting-dilution analyses

Spleens were homogenized in a tenBroek tissue disrupter. Red blood cells were lysed by incubating tissue homogenate in 8.3 g/L ammonium chloride for 10 minutes at room temperature with shaking. Cells were filtered through 40-micron mesh to reduce clumping. Frequencies of cells harboring MHV68 genomes were determined using a limiting-dilution, nested PCR analysis as previously described (44). Frequencies of latently-infected cells capable of reactivating were determined using a limiting-dilution analysis for cytopathic effect induced on an indicator MEF monolayer as previously described (44).

### Antibodies, tetramers, and treatments

CD19-BV650 (6D5), IgM-BV421 (RMM-1), CD38- Pacific Blue (90), and purified B220 (RA3-6B2) were purchased from Biolegend (San Diego, CA). CD3e-PCPCy5.5 (145-2c11), CD8*α*-PCPCy5.5 (53-6.7), CD4-AF700 (RMA-5), B220-AF700 (RA3-6B2), GL7-eF660 (GL-7), CD38-PE-Cy7 (90), CD138-BV711 (281-2), NK1.1-PCPCy5.5 (PK136), and IgD-PE (11-26c) were purchased from eBioscience. Other antibodies include mouse anti-p53 Alexa fluor 647-conjugate (Cell Signaling), goat anti-GFP (Rockland), and donkey anti-goat alexa fluor 488-conjugate (Invitrogen). MHV68-specific MHC class I tetramers were generated by the NIH Tetramer Core. As a positive control for p53 induction, bulk splenocytes were treated with 10 grays of gamma radiation by exposure to a cesium-137 source. Following exposure, cells were allowed to recover at 37°C in a tissue-culture incubator for 1 hour prior to staining for flow cytometry. Annexin V-PE was added along with antibodies to evaluate cell death.

### Flow cytometry

3 ⨯10^6^ cells were washed with FACS buffer (0.2% BSA, 1 mM in PBS) before blocking with Fc block (Invitrogen) and incubation with eF780 live/dead viability stain (eBioscience) for 10 minutes at 4°C. Surface staining was then performed with antibodies diluted at 1:300 for 30 minutes incubation time at 4°C. For intracellular stains, cells were fixed and permeabilized using a FoxP3 staining kit (eBioscience) following the manufacturers guidelines. Samples were analyzed using an LSRFortessa (Becton Dickinson). Data were analyzed using FlowJo (10.4.2) software.

### Immunohistology and microscopy

Tissues were fixed in 4% formaldehyde prior to paraffin embedding and cut into 5 µm sections. For H&E, sections were stained with hematoxylin and eosin as previously described (78). Tissue sections were imaged on an EVOS digital microscope with 10X or 40X objectives (Thermo). For immunohistochemical analyses, the sections were deparafinized, rehydrated, and incubated in citrate buffer for antigen retrieval before subject to staining with Vectastain Elite ABC kit (Vector) using a chromogenic reporter, DAB (Dako). Sections were counterstained with hematoxylin. Anti-fade mounting medium (Vector) was used prior to examining tissue sections on an Eclipse T5100 microscope (Nikon).

### Real-time PCR and genotyping

Lungs were harvested from mice infected with H2B-YFP MHV68 on day 7 post-infection and homogenized using 0.5 mm silica beads a Minibeadbeater (BioSpec) as previously described (44). DNA was extracted from homogenized tissue using a DNeasy kit (Qiagen) according to the manufacturer’s instructions followed by treatment with RNAse I (Qiagen) for 30 minutes at 37°C. Quantitative PCR was performed on 500 ng of DNA with primers forward corresponding to the MHV68 ORF59 genomic locus (forward: 5’-ATG-CAG-ACC-TTC-CAG-CTT-GAC-3’; reverse: 5’-CTC-TTC-CAA-GGG-AGC-TTG-CG-3’). Cycling parameters were 95°C for 30 seconds, 55°C for 30 seconds, and then 72°C for 30 seconds for 40 cycles on an ABIsteponeplus apparatus (ABI). Genotyping of mice was performed using a mouse genotyping kit (Kapa Biosystems). Briefly, DNA was extracted from mouse tail snips and amplified as previously described (79).

### Reverse transcriptase PCR

Splenocytes were isolated from H2B-YFP MHV68-infected or mock-infected mice at 16 days post-infection, and total RNA was extracted from the cells using Qiagen RNeasy Mini Kit (#74104). 1 ug RNA was used for cDNA synthesis (Invitrogen) prior to PCR analysis following manufacturer’s guidelines (Agilent). Gene-specific TaqMan probes (Applied Biosystems) for Mdm2 (Mm01233136_m1), Cdkn1a (Mm00432448_m1) and β-actin (Mm00607939_s1) were utilized for quantification. Reactions were performed in an Applied Biosystems StepOnePlus PCR system with cycling conditions of 10 min at 95°C followed by 40 cycles of 15 s at 95°C and 1 min at 60°C. Biological triplicate samples were analyzed in technical triplicate using the ΔΔCT method with β-actin as the cellular housekeeping transcript control.

### c-Myc translocation PCR

High molecular weight genomic DNA was extracted from P3×63Ag cells, B cells, bulk splenocytes or 3T12 fibroblasts using proteinase K and phenol (80). For positive translocation control, primary B cells from spleens of p53^-/-^ mice were enriched by immunomagnetic depletion of CD3^+^ cells (Miltenyi Biotech) and stimulated with 25 μg/ml LPS and 5 ng/ml IL-4 in RPMI 1640 medium for four days to induce class-switch recombination (54). After stimulation, live B cells were isolated with Lympholyte-M (Cedarlane) according to manufacturer’s instruction. Two-rounds of translocation PCR was performed using GoTaq G2 Green Master Mix (Promega). In the first round PCR, genomic DNA corresponding to 10^5^ cells was used in each reaction with the following primers: for derChr12, Ig-R1 5’-TCATCCCGAACCATCTCAACCAG-3’ and Myc-R1 5’-GACACCTCCCTTCTACACTCTAAACCG-3’; for derChr15, Ig-F1 5’- GCACAGCTGAGCTGAGATGG-3’ and Myc-F1 5’-TCCTGCATAGACCTCATCTGCG-3’. In the second round PCR, a 1 μl sample of the first round PCR was added to each reaction with the following primers: for derChr12, Ig-R2 5’-TCTGAGCCTAGTTCAACCTGGC-3’ and Myc-R2 5’- CACTGCACCAGAGACCCTGC-3’; for derChr15, Ig-F2 5’-GTGGGCTTCTCTGAGTGCTTCT-3’ and Myc-F2 5’-CGGTTGATCACCCTCTATCACTCC-3’. For loading control PCR, 250 ng of high molecular weight DNA was amplified with primers Myc-F1 and Myc-R1. PCR conditions were 94°C for 2 min, followed by 30 cycles of 94°C for 30 s, 60°C for 30 s and 72°C for 3 min, and final extension of 72°C for 1 min. Translocation positive bands were re-amplified from the corresponding first round PCR in second round PCR reactions and purified with QIAquick Gel Extraction Kit (Qiagen) for DNA sequencing.

### EdU incorporation assays

Mice were injected with 100 μg of 5-ethynyl-2’deoxyuridine (EdU, Invitrogen) 2 hours prior to splenocyte harvest. Cells were fixed with 4% paraformaldehyde in PBS and Click-iT chemistry (BD Pharmagen) performed according to the manufacturer’s instructions to fluorescently mark EdU+ cells.

### Enzyme-linked immunosorbent assays

Viral antigen was prepared by infecting 3T12 fibroblasts at an MOI of 0.5 PFU/cell for 96 hours. Cells were washed and fixed in 1% PFA. Viral antigen was used to coat high-binding plates (Denville) overnight at 4°C. After blocking in 5% FBS in 1× PBS, serially diluted serum was incubated on the plate, followed by incubation with HRP- conjugated IgM or IgG Ab (Southern Biotech). SureBlue substrate (KPL) was added to detect the Ag-specific Abs. The wells were read at 450 nm on an FLUOstar Omega plate reader (BMG Labtech).

---

**Supplemental Figure 1.**
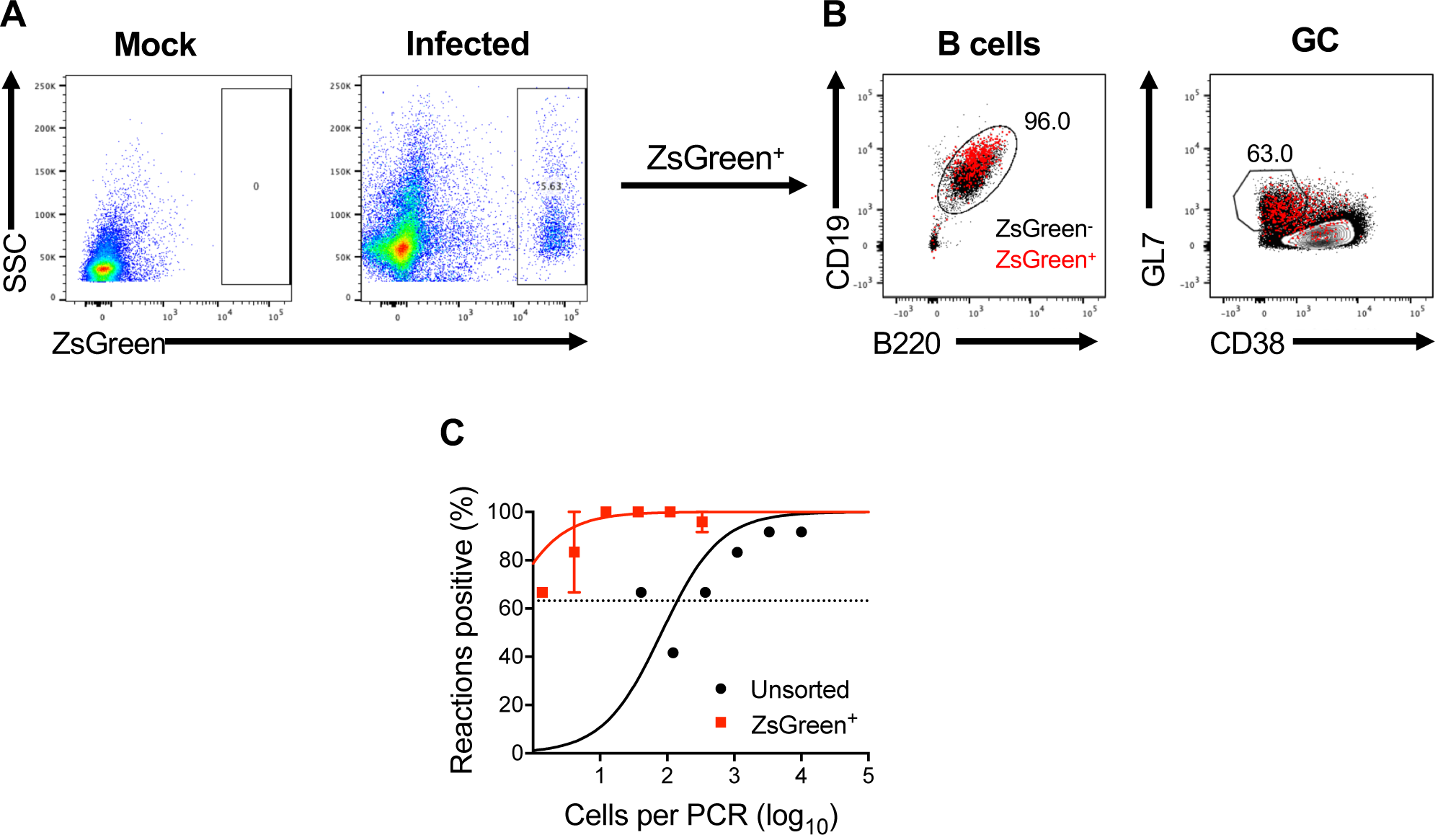
ZsGreen^+^ cells are infected with MHV68-Cre. (A-C) ZsGreen mice were infected intranasally with MHV68-Cre (10^4^ PFU). Mice were sacrificed and splenocytes harvested on day 16 post-infection. (A) Representative flow plots for ZsGreen expression. (B) Harvested splenocytes were stained with cell specific markers for the indicated cell types. B cells were gated as CD19^+^/B220^+^, GC B cells as GL7^+^/CD38^lo^. ZsGreen^+^ B cell populations indicated in red, ZsGreen^-^ B cell populations indicated in black. Population frequencies apply to ZsGreen^+^ cells. (C) ZsGreen^+^ cells were sorted and limiting dilution PCR was performed to quantify proportion of splenocytes harboring latent viral genomes.

**Supplemental Figure 2.**
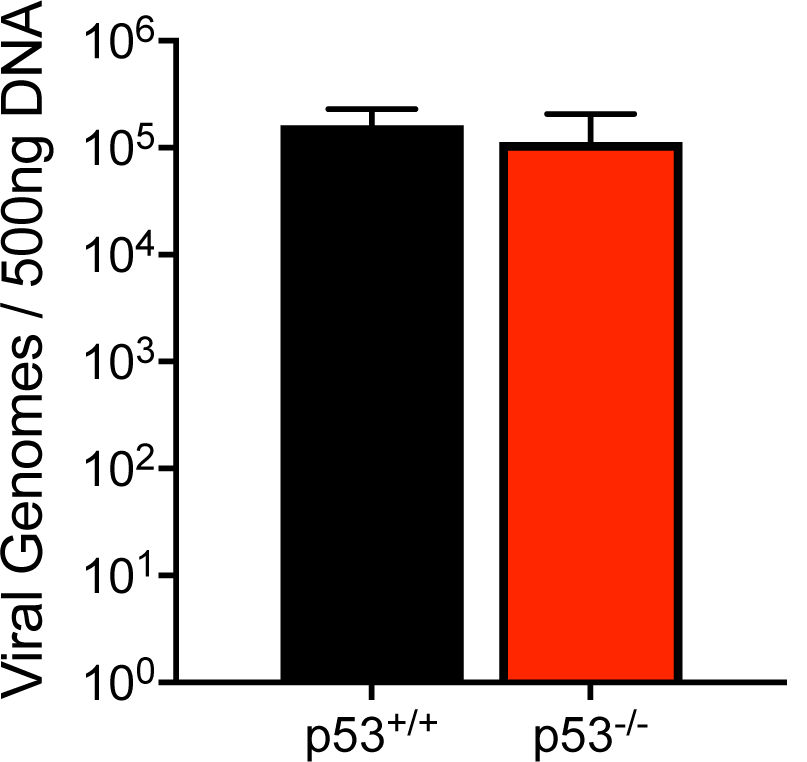
p53 does not affect MHV68 acute replication. p53^+/+^ or p53^-/-^ mice (n=3) were intranasally inoculated with 10^4^ PFU of H2B-YFP-expressing MHV68. Animals were sacrificed 7 days post-infection and DNA was isolated from lungs for quantitative PCR to detect viral genomes. Results are means of 3 samples. Error bars represent standard deviation.

**Supplemental Figure 3.**
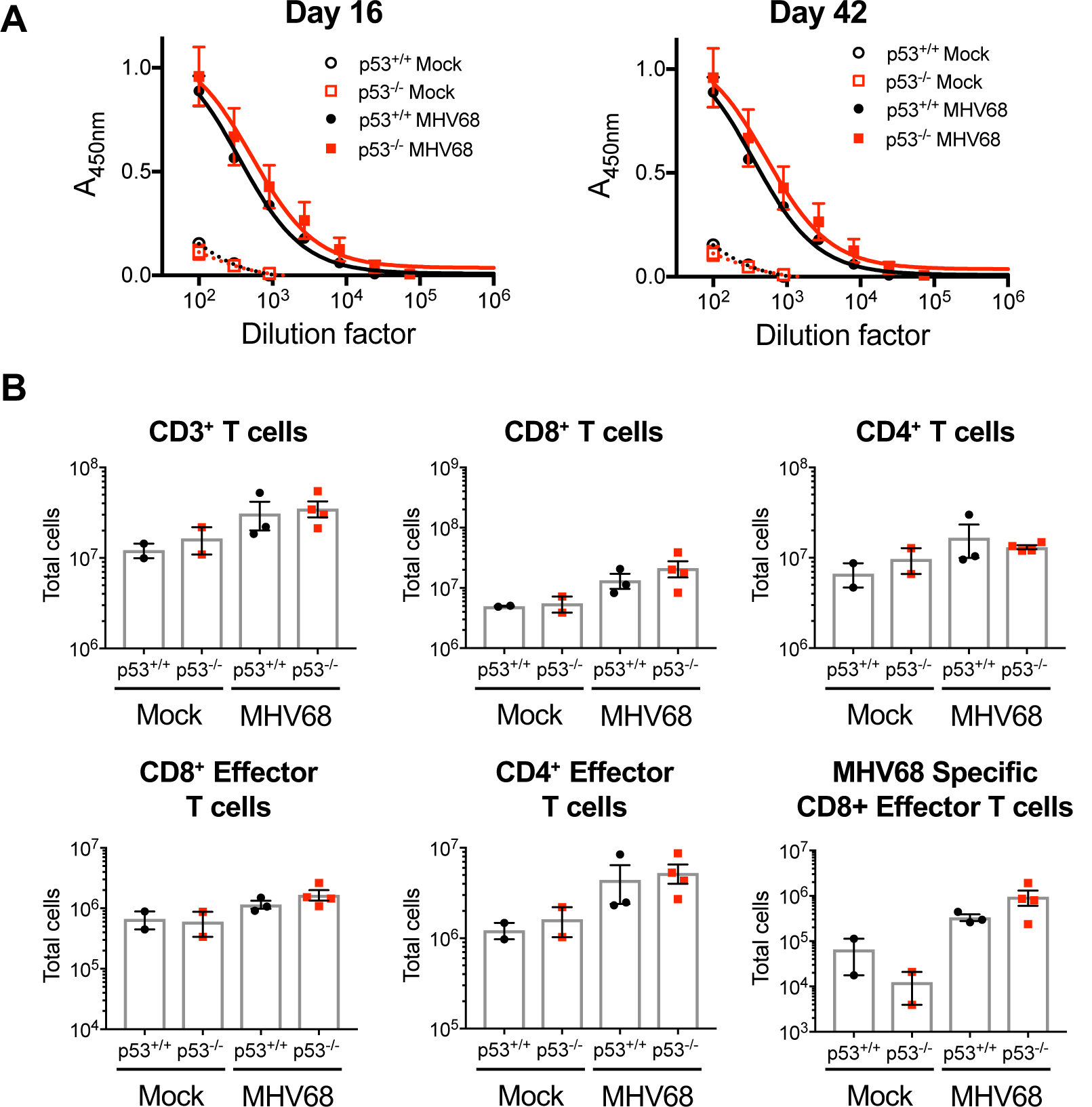
p53 is not required for virus-specific adaptive immunity. (A-B) p53^+/+^ or p53^-/-^ mice (n=3) were intranasally inoculated with 10^4^ PFU of H2B-YFP-expressing MHV68. (A) Blood samples were harvested from the submandibular vein at days 16 and 42 post-infection. ELISAs were performed on serum using MHV68 antigen. (B) On day 42 post-infection mice were sacrificed, splenocytes harvested and stained with specific antibodies against CD3, CD8, CD19, and PE-conjugated MHCI tetramers specific for MHV68 ORF6. Flow cytometry was performed to quantify total T cells (CD19^-^/CD3^+^), CD8^+^ T cells, CD4^+^ T cells, effector T cells (CD62L^-^/CD44^+^) and MHV68-specific effector T cells (tet^+^). Representative data of two independent experiments. Results are expressed as means and error bars represent standard error of mean. Mann-Whitney unpaired t test.

**Supplemental Figure 4.**
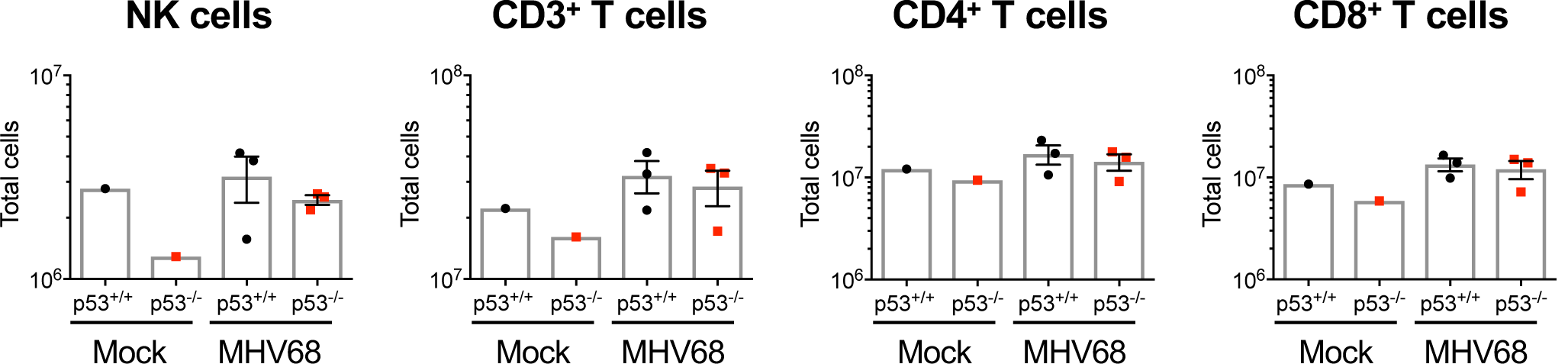
p53 expression is not required in NK and T cell populations during MHV68 infection. p53^+/+^ or p53^-/-^ littermates were infected intranasally with 10^4^ PFU of H2B-YFP MHV68 (n=3). Mice were sacrificed on day 16 post-infection and splenocytes were harvested. Harvested splenocytes were stained with cell specific markers for the indicated cell types. T cells were gated as CD3^+^, T cell subtypes identified by CD8 or CD4 expression. Natural killer cells were gated as CD3^lo^/NK1.1^+^. Live cells were identified with Viability Dye eFluor 780 (eBioscience). Each dot represents one mouse. Results are expressed as means and error bars represent standard error of mean.

**Supplemental Figure 5.**
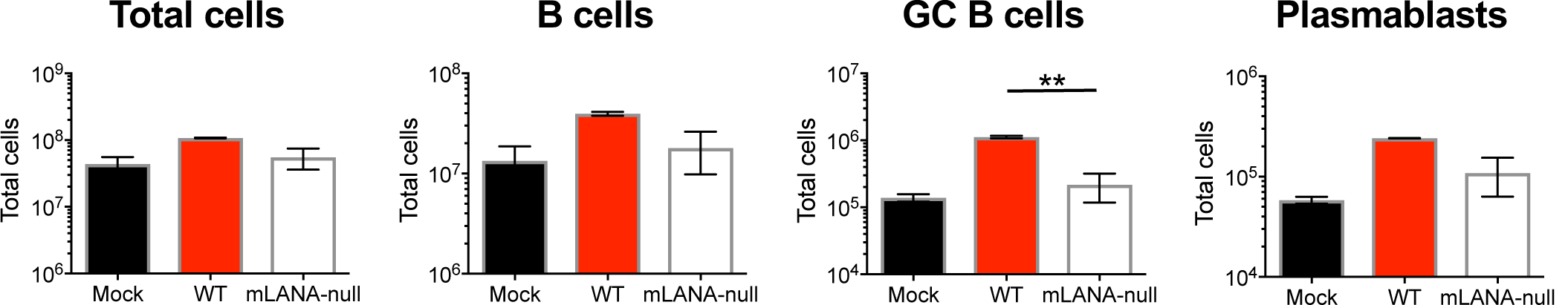
Latency establishment is required for splenic B cell expansion. p53^-/-^ mice were infected intranasally with 10^4^ PFU of H2B-YFP MHV68 or mLANA-null MHV68 (n=3). Mice were sacrificed on day 16 post-infection and splenocytes were harvested. Harvested splenocytes were stained with cell specific markers for the indicated cell types. B cells were gated as CD19^+^/B220^+^, GC B cells as GL7^+^/CD38^lo^, and plasmablasts as CD138^+^/B220^lo^. Live cells were identified with Viability Dye eFluor 780 (eBioscience). Results are means +/- SEM. Mann-Whitney unpaired t test * p<0.005

**Supplemental Figure 6.**
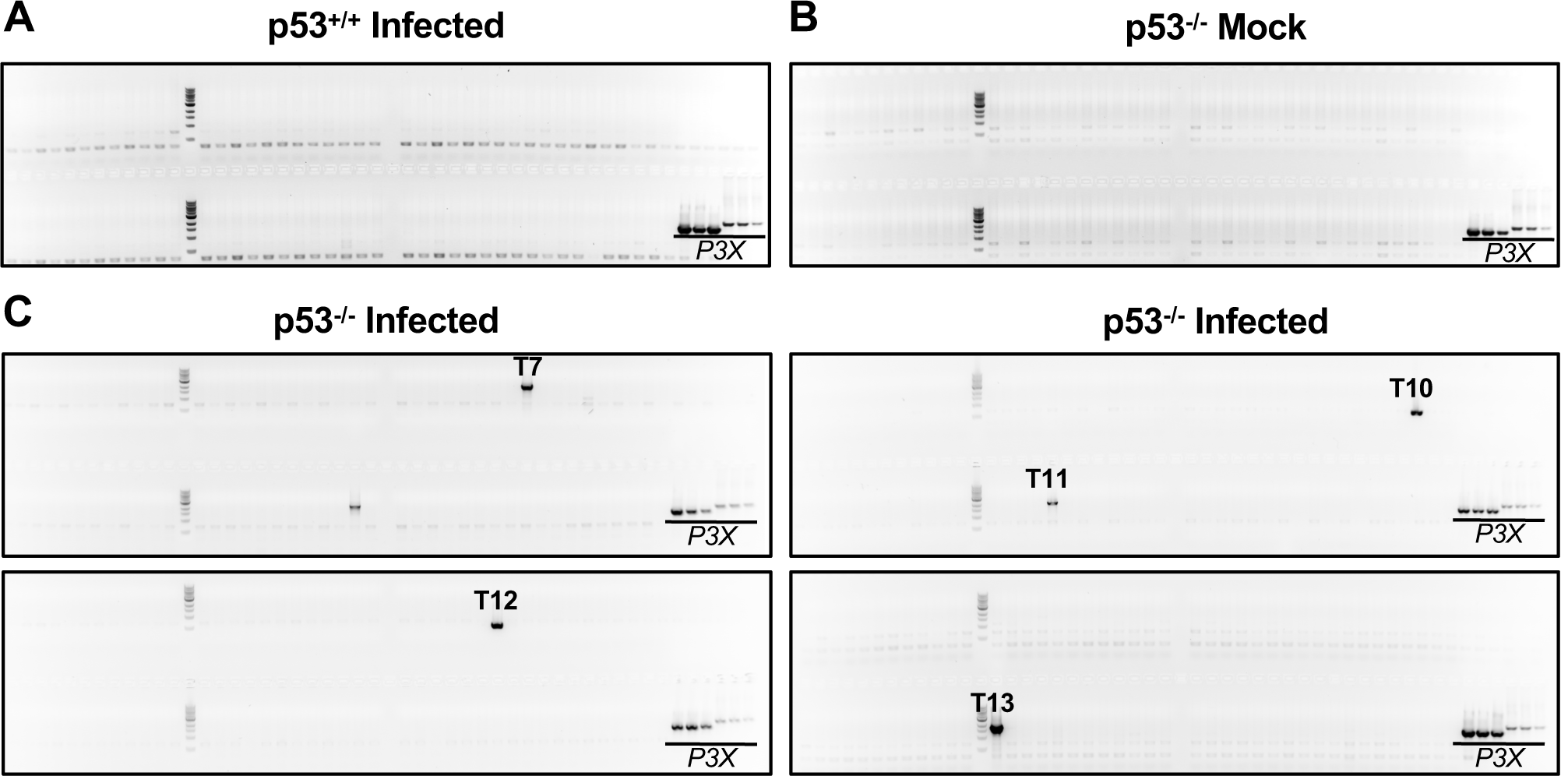
Infection of p53^-/-^ mice causes genomic instability. (A-C) p53^+/+^ or p53^-/-^ littermates were infected intranasally with 10^4^ PFU of H2B-YFP MHV68 (n=5). Mice were sacrificed on day 16 post-infection and splenocytes were harvested. B cells were enriched by MACS isolation and pooled, and chromosomal DNA extracted. DNA from 10^5^ cells were added in each translocation PCR. Positive bands were cut and purified for DNA sequencing. P3X cells served as a positive control for translocation. (A) Representative translocation PCR of p53^+/+^ infected animals. (B) Representative translocation PCR of p53^-/-^ mock animals. (C) Translocation PCR from infected p53^-/-^ mice. Labels refer to nucleotide sequences in Table 2.

**Supplemental Figure 7:**
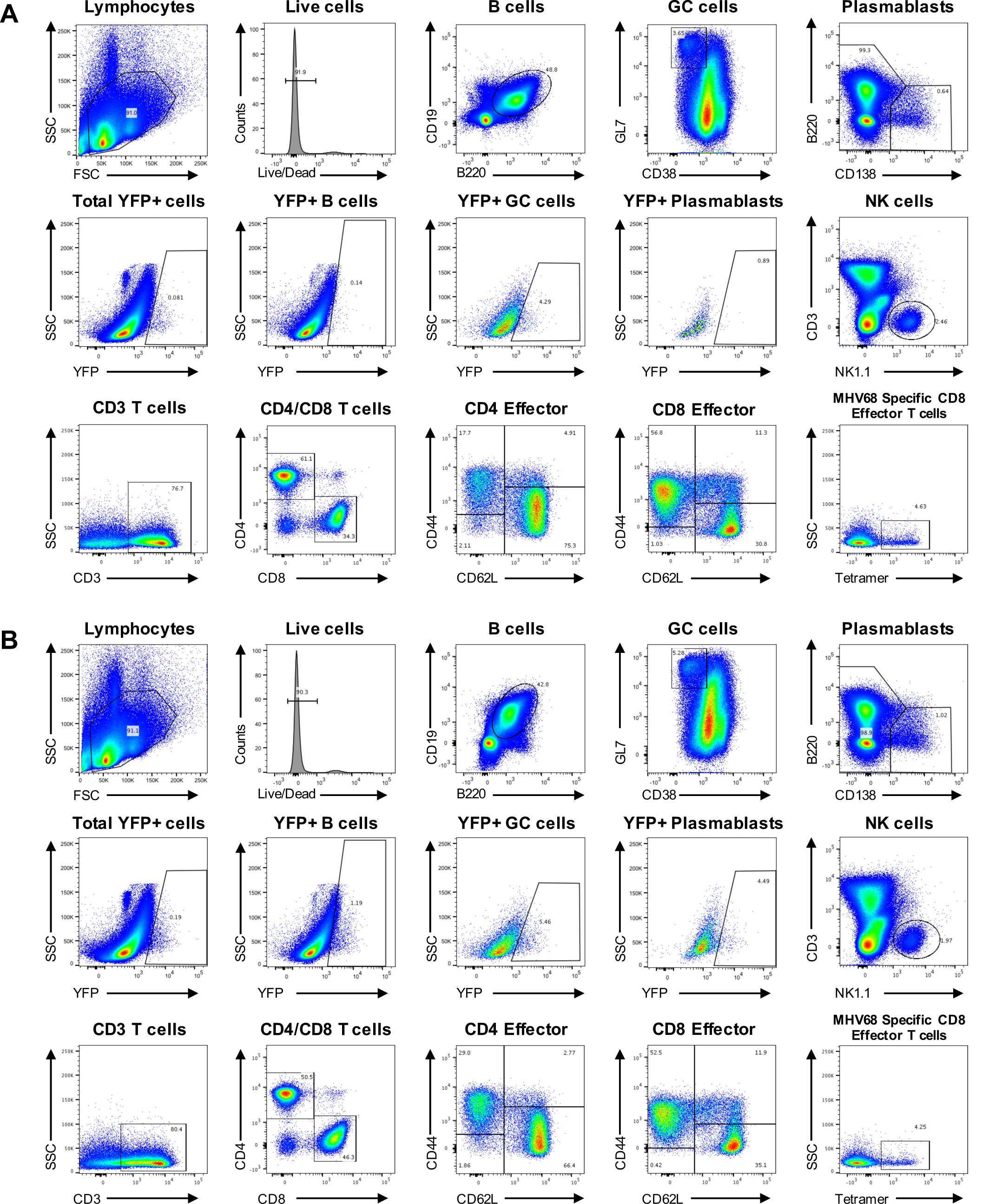
Representative flow plots for cell phenotyping experiments. (A-B) p53^+/+^ or p53^-/-^ littermates were infected intranasally with 10^4^ PFU of H2B-YFP MHV68 (n=3). Mice were sacrificed on day 16 post-infection and splenocytes were harvested. Harvested splenocytes were stained with cell specific markers for the indicated cell types. B cells were gated as CD19^+^/B220^+^, GC B cells as GL7^+^/CD38^lo^, plasmablasts as CD138^+^/B220^lo^ cells. T cells (CD3^+^), CD8^+^ T cells, CD4^+^ T cells, effector T cells (CD62L^-^/CD44^+^) and MHV68-specific effector T cells (tet^+^). NK cells were gated as CD3^lo^/NK1.1^+^. (A) p53^+/+^ mice. (B) p53^-/-^ mice.

**Supplemental Figure 8:**
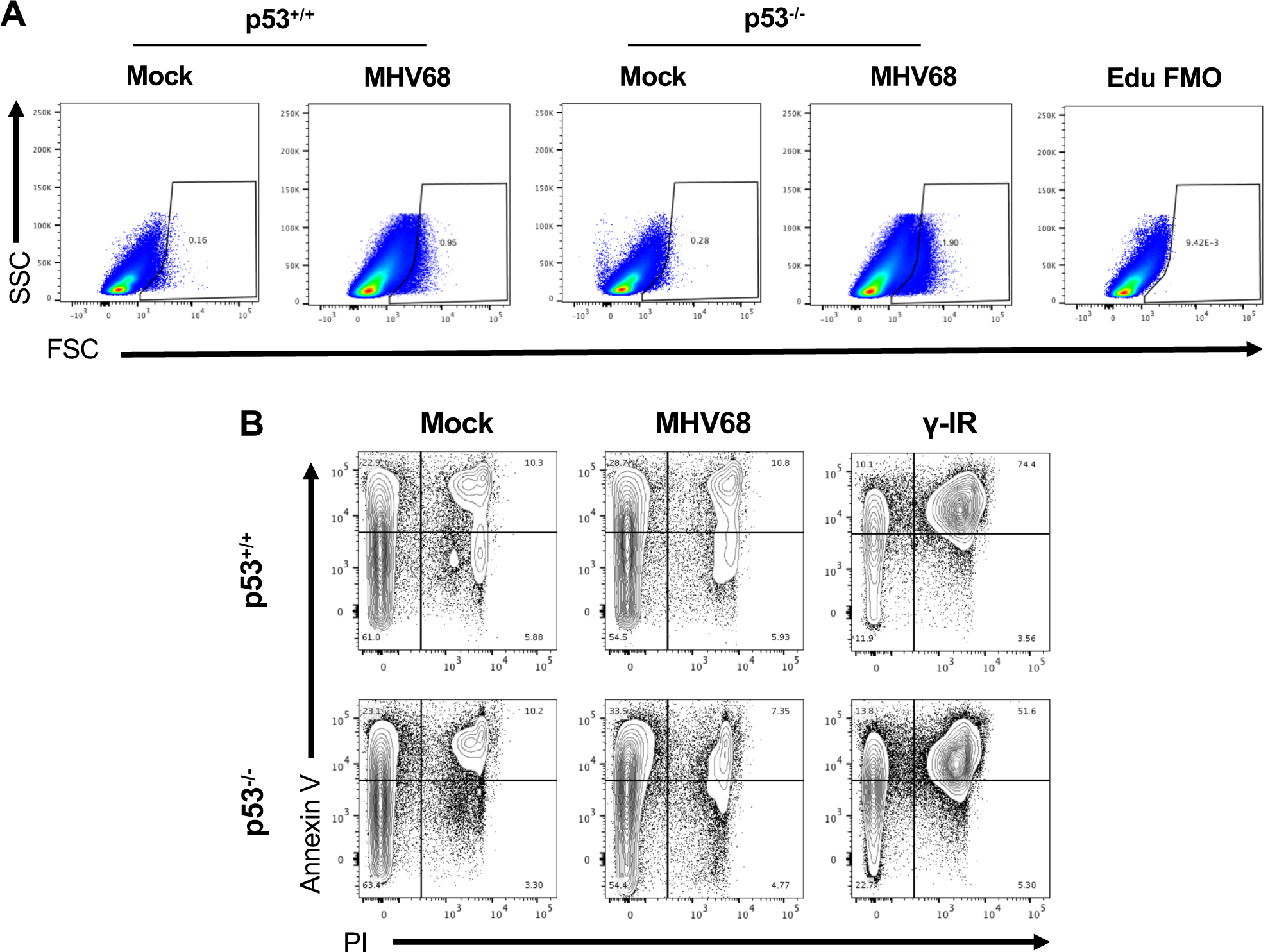
Representative flow plots for EdU incorporation and Annexin V assay. (A) Representative flow plots from experiments described in Figure 4A. (B) Representative flow plots from experiments described in Figure 4B.

**Supplemental Figure 9:**
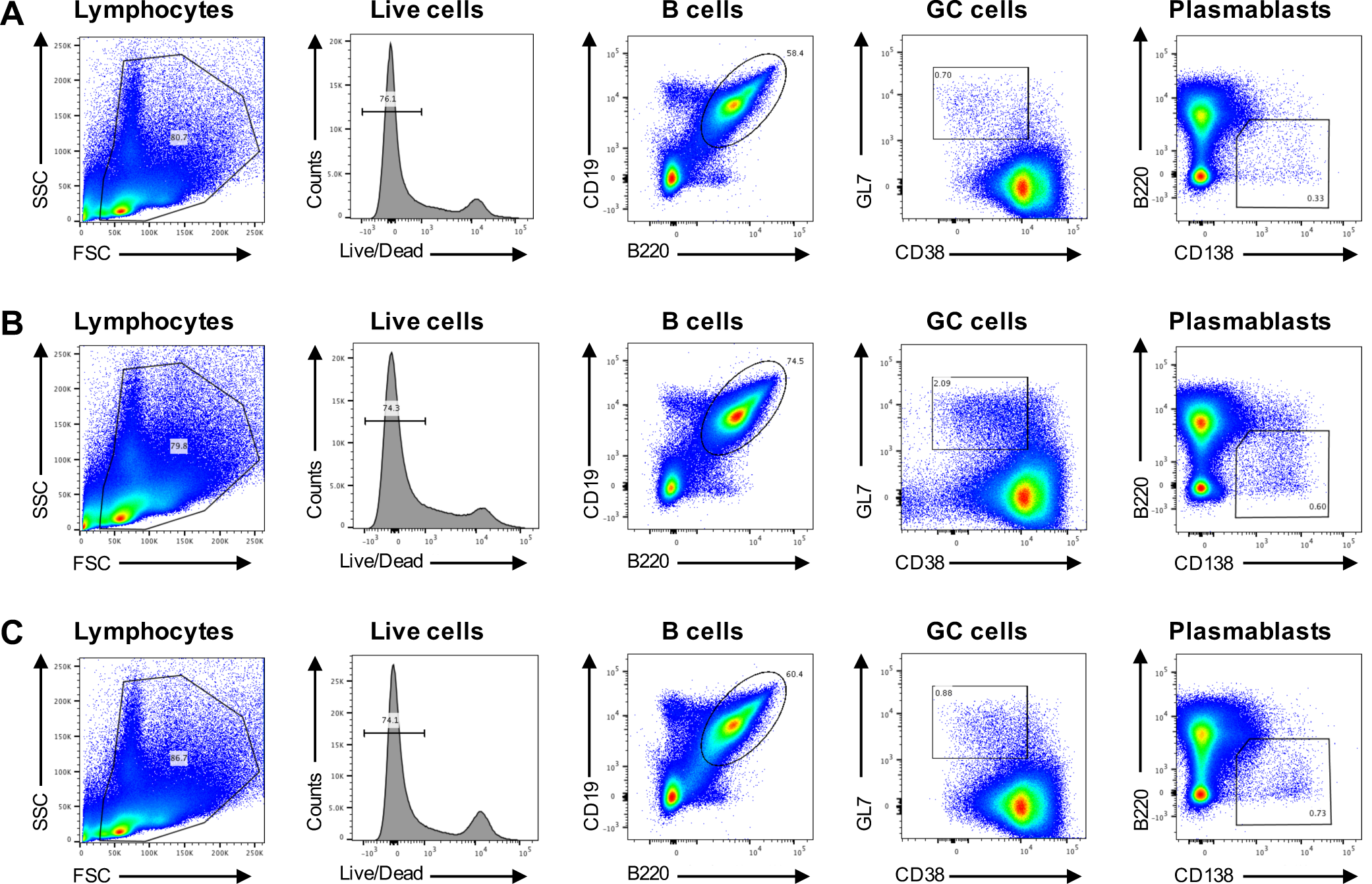
Representative flow plots for mLANA-null infections of p53^-/-^ mice. (A-C) Representative plots from experiments described in Supplemental Figure 4. (A) Mock infected mice. (B) H2B-YFP MHV68 infected mice. (C) mLANA-null MHV68 Infected mice.

